# Fine Mapping and Candidate Gene Analysis of Dravet Syndrome Modifier Loci on Mouse Chromosomes 7 and 8

**DOI:** 10.1101/2024.04.15.589561

**Authors:** Nicole A. Hawkins, Nathan Speakes, Jennifer A. Kearney

## Abstract

Dravet syndrome is a developmental and epileptic encephalopathy (DEE) characterized by intractable seizures, comorbidities related to developmental, cognitive, and motor delays, and a high mortality burden due to sudden unexpected death in epilepsy (SUDEP). Most Dravet syndrome cases are attributed to *SCN1A* haploinsufficiency, with genetic modifiers and environmental factors influencing disease severity. Mouse models with heterozygous deletion of *Scn1a* recapitulate key features of Dravet syndrome, including seizures and premature mortality; however, severity varies depending on genetic background. Here, we refined two Dravet survival modifier (*Dsm*) loci, *Dsm2* on chromosome 7 and *Dsm3* on chromosome 8, using interval-specific congenic (ISC) mapping. *Dsm2* was complex and encompassed at least two separate loci, while *Dsm3* was refined to a single locus. Candidate modifier genes within these refined loci were prioritized based on brain expression, strain-dependent differences, and biological relevance to seizures or epilepsy. High priority candidate genes for *Dsm2* include *Nav2, Ptpn5, Ldha, Dbx1, Prmt3* and *Slc6a5*, while *Dsm3* has a single high priority candidate, *Psd3*. This study underscores the complex genetic architecture underlying Dravet syndrome and provides insights into potential modifier genes that could influence disease severity and serve as novel therapeutic targets.

## INTRODUCTION

Dravet syndrome is a developmental and epileptic encephalopathy (DEE) that arises during infancy (Dravet 2011). Initial seizures often occur in the context of a fever; however, the syndrome evolves to include afebrile seizure types. Seizures are often prolonged and respond poorly to conventional anticonvulsants (Dravet and Oguni 2013). Beyond seizures, individuals with Dravet syndrome exhibit several comorbidities, including developmental delay, intellectual disability, motor impairment, and behavioral and/or psychiatric issues, each impacting quality of life (Darra et al. 2019; Feng et al. 2024; Villas et al. 2017). Dravet syndrome is associated with a significantly elevated mortality risk, ranging from 4.4-17.5%, typically occurring during childhood. These premature deaths primarily result from sudden unexplained death in epilepsy (SUDEP) (15-53% of cases) or status epilepticus (SE) (11.5-36% of cases) (Cooper et al. 2016; Dravet et al. 2005; Sakauchi et al. 2011a, b; Strzelczyk et al. 2023).

Over 80% of individuals with Dravet syndrome have *de novo* pathogenic variants in the *SCN1A* gene that encodes the NaV1.1 voltage-gated sodium channel (Depienne et al. 2009; Zuberi et al. 2011). *SCN1A* haploinsufficiency is the major mechanism underlying Dravet syndrome (Gallagher et al. 2024). Despite a shared genetic basis, there is a range of clinical severity among individuals with *SCN1A* haploinsufficiency (Depienne et al. 2010; Depienne et al. 2009; Goldberg-Stern et al. 2014; Guerrini et al. 2010; Nabbout et al. 2003; Osaka et al. 2007; Yordanova et al. 2011; Yu et al. 2010). This suggests that clinical phenotypes are influenced by other factors, including genetic modifiers and environmental factors (de Lange et al. 2020; Hammer et al. 2017; Kearney 2011).

Dravet syndrome can be modeled in mice by heterozygous deletion of *Scn1a*. Several mouse models with varying deletions have been generated and share similar phenotypes, including spontaneous and hyperthermia-induced seizures (Miller et al. 2014; Ogiwara et al. 2007; Yu et al. 2006). Similarly, heterozygous knock-in of loss-of-function missense variants can also recapitulate core Dravet syndrome features (Ricobaraza et al. 2019). Both knockout and knock-in models have a high risk of sudden unexpected death following a seizure in otherwise healthy animals (SUDEP-like) (Kalume et al. 2013; Miller et al. 2014; Ricobaraza et al. 2019; Yu et al. 2006). In the knockout models, the highest incidence of SUDEP-like deaths occurs early in life, with approximately 50% mortality by one month of age (Kalume et al. 2013; Miller et al. 2014).

A consistent feature among Dravet mouse models is variable penetrance and expressivity of Dravet-like phenotypes dependent on genetic strain background (Miller et al. 2014; Rubinstein et al. 2015; Yu et al. 2006). We generated the *Scn1a*^*tmKea*^ model with heterozygous deletion of the first coding exon (abbreviated *Scn1a*^*+/-*^) (Miller et al. 2014). Phenotype severity was highly dependent on strain background. On the 129S6/SvEvTac (129) strain, *Scn1a*^*+/-*^ mice did not develop epilepsy and lived a normal lifespan. In contrast, when crossed with C57BL/6J (B6) mice, the resulting [B6 x 129]F1.*Scn1a*^*+/-*^ mice recapitulated core features of Dravet syndrome, including spontaneous seizures and epilepsy, and a high incidence of SUDEP-like death in the first month of life (Miller et al. 2014). We previously performed genetic mapping to identify Dravet survival modifier (*Dsm*) loci influencing strain-dependent survival of *Scn1a*^*+/-*^ mice. Five *Dsm* loci were localized to mouse chromosomes 5, 7, 8 and 11 (Miller et al. 2014). *Dsm1* and *Dsm4* mapped to an overlapping region on chromosome 5 in both 129 and B6 backcrosses (Miller 2014). Further fine mapping and candidate gene analyses of *Dsm1/4* locus on the chromosome 5 identified and validated *Gabra2* as a modifier gene (Hawkins et al. 2021; Hawkins et al. 2016; Nomura et al. 2019). In the current study, we used a similar interval-specific congenic (ISC) mapping approach to refine *Dsm2* and *Dsm3* on chromosomes 7 and 8, respectively. In addition, we performed candidate gene analysis within the refined intervals to nominate putative modifier genes that may influence strain-dependent survival of *Scn1a*^*+/-*^ Dravet mice.

## METHODS

### Mice (NCBI Taxon ID 10090)

*Scn1a*^*tm1Kea*^ mice (RRID:MMRRC 037107-JAX) are maintained as an isogenic strain by continued backcrossing to 129S6/SvEvTac inbred mice (129SVE, Taconic Biosciences, Germantown, NY, USA), and abbreviated herein as 129.*Scn1a*^*+/-*^ (Miller et al. 2014). C57BL/6J inbred mice were obtained from the Jackson Laboratory (Bar Harbor, ME, USA; RRID:IMSR_JAX:000664). Mice were maintained in a Specific Pathogen Free (SPF) barrier facility with a 14:10 light:dark cycle and access to food and water *ad libitum*. Animal care and experimental procedures were approved by the Northwestern University Animal Care and Use Committee in accordance with the National Institutes of Health Guide for the Care and Use of Laboratory Animals. The principles outlined in the ARRIVE (Animal Research: Reporting of in vivo Experiments) guidelines were considered when planning experiments (Percie du Sert et al. 2020).

### Interval Specific Congenic (ISC) Lines

ISC lines were generated by crossing 129SvEvTac (129) males with B6 females, and then successively crossing to C57BL/6J (B6) to generate congenic lines with 129-derived alleles on the chromosome of interest on a B6 background. Separate series of ISC strains were developed for *Dsm2* (designated as B6.129-ISC7A through B6.129-ISC7H) on chromosome 7, and for *Dsm3* (designated as B6.129-ISC8A through B6.129-ISC8G) on chromosome 8. At each generation mice were genotyped for markers in the region of interest and mice retaining 129 alleles were propagated. Whole genome and selective genotyping were performed at generations N4 and N7 to select breeders with low percentages of 129 in the rest of the genome. All lines were crossed to B6 for ≥N8 generations prior to any experiments.

### Genotyping

DNA was prepared from tail biopsies obtained at approximately 2-weeks of age (Gentra Puregene Mouse Tail Kit, Qiagen, Valencia, CA, USA). *Scn1a* genotype was determined as previously described (Kearney et al. 2022). Microsatellite genotypes were determined by analysis of PCR products on 7% nondenaturing polyacrylamide gels. Breakpoints for ISC lines were refined by SNP genotyping using the mini Mouse Universal Genotyping Array (miniMUGA) (Transnetyx, Cordova, TN).

### Phenotyping

Female B6.129-ISC7 or B6.129-ISC8 mice were bred with heterozygous 129.*Scn1a*^*+/-*^ males to generate F1 offspring carrying homozygous 129/129 alleles or heterozygous 129/B6 alleles in the *Dsm2* or *Dsm3* interval, respectively. This breeding scheme neutralizes confounding parent-of-origin effects for imprinted genes, a number of which reside on mouse chromosome 7 (Morison and Reeve 1998). At P12-14, mice were ear-tagged, tail biopsied and genotyped. Mice were weaned at P19-21 into standard vivarium cages containing 4-5 mice of the same sex/age and monitored for survival to 8 weeks age. Over the monitoring period, mice were checked daily for general health. Any visibly unhealthy mouse (e.g., underweight, dehydrated, poorly groomed, or immobile) was euthanized and excluded (rare) because the focus of the study was sudden and unexpected death in otherwise healthy appearing *Scn1a*^*+/-*^ mice (SUDEP-like). Subjects surviving beyond P56 were censored for survival analysis because they reached the predetermined end of the study period. *Dsm2* and *Dsm3* were analyzed separately with their respective ISC strains. Survival was compared between groups using Kaplan–Meier analysis with hazard ratios and *P*-values determined by LogRank Mantel–Cox tests. Group sizes were determined by data simulations using data from our prior survival studies (Hawkins et al. 2016; Kearney et al. 2022; Miller et al. 2014).

### Candidate Gene Analysis

Candidate gene sets were extracted from the Mus musculus GRCm39 reference genome using the Ensembl BioMart tool (Martin et al. 2023). The subset of genes expressed in brain were defined using the MGI Gene eXpression Database (Baldarelli et al. 2021). Differential expression of candidate genes between 129 and B6 or F1 was assessed using two RNA-Seq datasets that we previously reported: (1) B6 and 129 forebrain (Hawkins et al. 2016); and (2) F1 and 129 hippocampus (Hawkins et al. 2019) (NCBI GEO GSE112627). Coding sequence changes between strains were derived from our previously reported whole genome re-sequencing of 129S6/SvEvTac (NCBI SRA PRJNA817075) aligned to the C57BL/6J reference sequence (Kearney et al. 2022). Variants classified as missense, indel, or protein truncating were retained, and variants are reported with dbSNP Reference SNP (rs) accession numbers whenever possible. The predicted effect of missense variants on the protein encoded by the canonical transcript was assessed with SIFT and PolyPhen2 (Table S1) (Adzhubei et al. 2010; Vann et al. 2018).

## RESULTS

We previously reported low resolution mapping of *Dsm* loci that influenced strain-dependent survival of *Scn1a*^*+/-*^ mice (Miller et al. 2014). In that report, we demonstrated that 129 alleles conferred risk at *Dsm2* on chromosome 7, whereas 129 alleles were protective at *Dsm3* on chromosome 8 (Miller et al. 2014). To refine the map intervals for these loci, we generated ISC lines carrying 129-derived alleles in the regions of interest on an otherwise B6 background. Separate series of lines were generated and analyzed for *Dsm2* on chromosome 7 (designated ISC7-letter) and *Dsm3* on chromosome 8 (designated ISC8-letter). For each line, ISC females were crossed to 129.*Scn1a*^*+/-*^ males and survival was monitored to 8 weeks of age. Survival was then compared between offspring carrying homozyogous (129/129) alleles in the ISC region or heterozygous (129/B6) F1 controls. Based on our initial mapping, we expected that 129 homozygosity would result in worse survival for modifier-containing intervals on chromosome 7 and improved survival for modifier-containing intervals on chromosome 8 (Kearney et al. 2022).

### Dsm2 Fine mapping and Candidate Gene Analysis

To map *Dsm2*, we used eight ISC lines carrying 129-derived alleles across the interval on chromosome 7 (Fig. 1A). The region on chromosome 7 is complex and likely has multiple contributing loci, similar to our previous report of *Dsm5* on mouse chromosome 11 (Kearney et al. 2022). Line ISC7-H confers significant risk, while line ISC7-C, confers moderate risk (Fig. 1B,C) (Table 1). Lines ISC7-F and ISC7-B also trend toward moderate risk. We interpret this as there being proximal and distal risk alleles that synergize in line ISC7-H (Fig. 1A). Potential protective contributions from distal regions of the mapped interval may attenuate risk in lines ISC7-C, ISC7-B, ISC7-G, and ISC7-A (Fig. 1A). The map intervals of interest for *Dsm2* are 25.3-34.1 Mb (*Dsm2a*) and 48.6-52.9 Mb (*Dsm2b*) on chromosome 7.

**Table 1.**
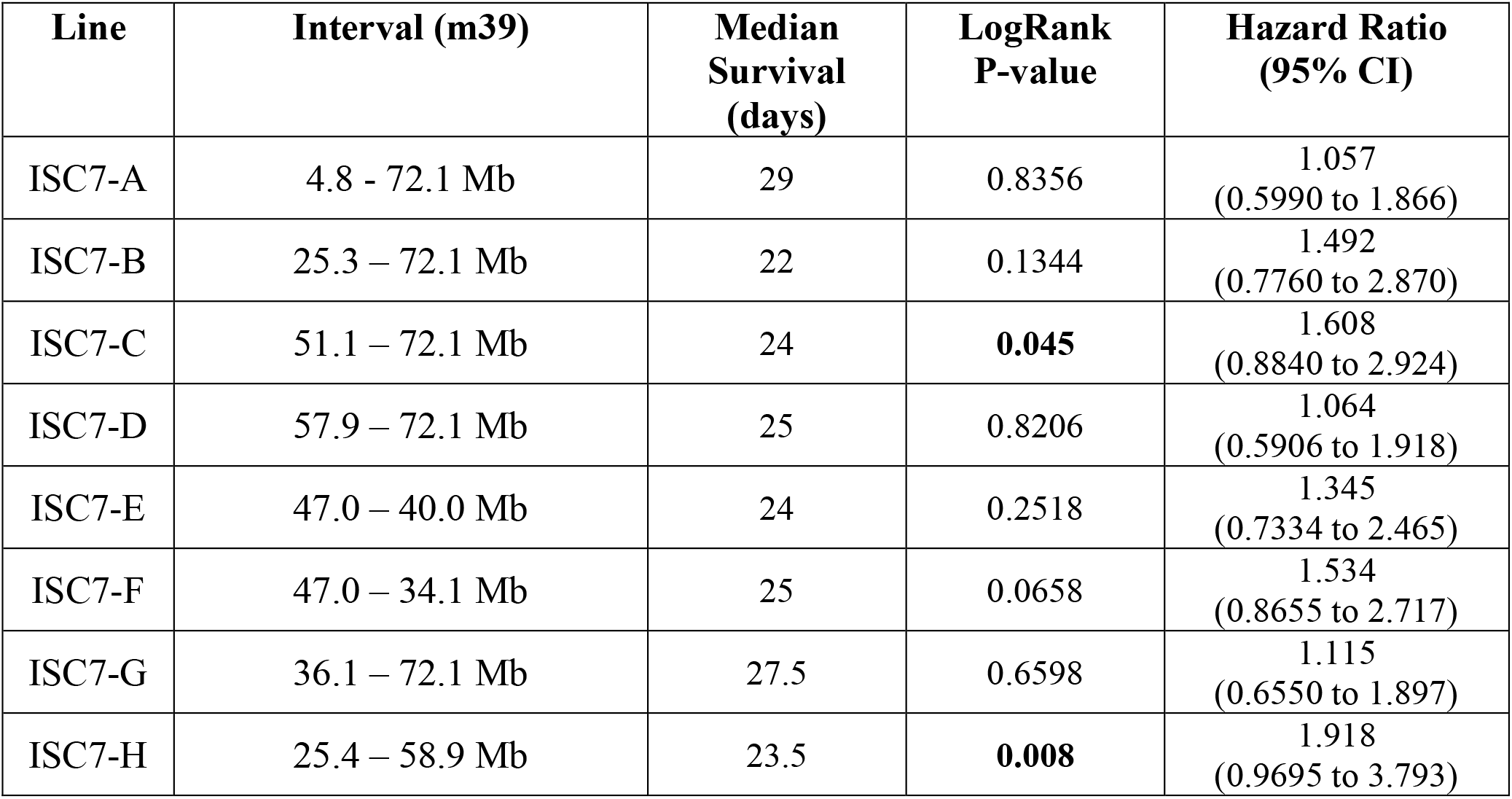
Interval specific congenic mapping of *Dsm2* on chromosome 7.

**Figure 1.**
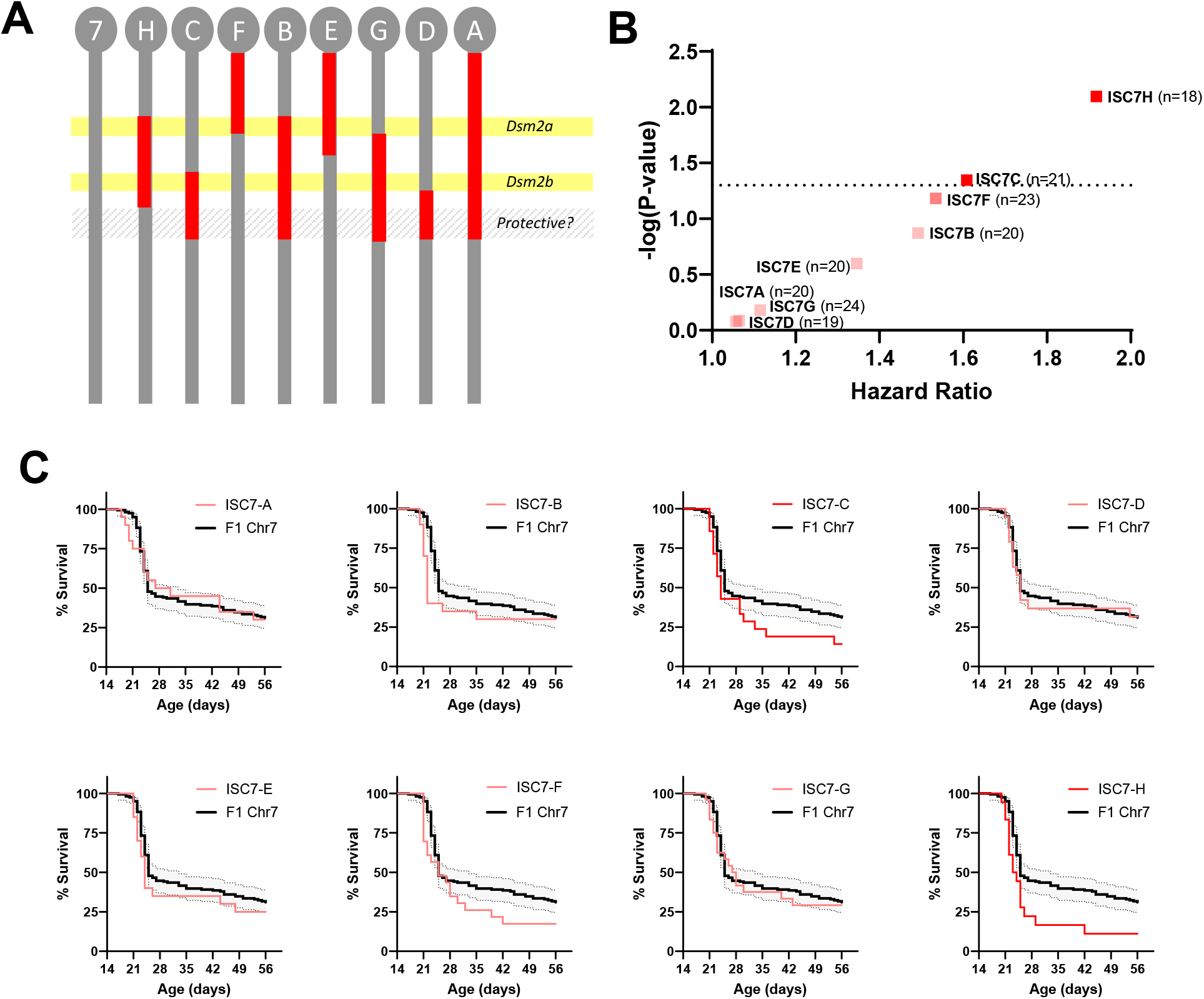
Fine mapping of *Dsm2* with ISC strains. **A)** *Dsm2* lines have varying 129-derived intervals (red) on a congenic B6 background (grey). B6.129-*Dsm2* ISC7 lines were crossed with 129.*Scn1a*^*+/*−^ mice and survival of *Scn1a*^*+/*−^ offspring was monitored to 8 weeks of age. **B)** Hazard ratios for all *Dsm2* ISC lines compared to F1 controls plotted against -log10(*P*-values) determined by Mantel-Cox LogRank test (n=18-24 per line). **C)** Kaplan Meier survival plots are shown for each ISC7 line (red) compared to F1.*Scn1a*^*+/-*^ controls (black). Shaded area represents 95% confidence interval for F1. *Scn1a*^*+/-*^ controls.

At the current resolution, the *Dsm2a* region contains 224 known protein coding genes and 36 noncoding RNA genes, and will require additional refinement to support candidate gene prioritization. In contrast, the *Dsm2b* interval is tractable and contains 51 known protein coding genes and 5 known noncoding RNA genes. Among the 51 protein coding genes in *Dsm2b*, 32 are expressed in brain (Table 2). We evaluated the brain expressed candidates for evidence of differential gene expression (DEG) in our existing mRNA-seq datasets (Hawkins et al. 2019; Miller et al. 2014). We found a total of five genes with evidence of differential expression (Table 2). There were three DEGs when comparing F1.*Scn1a*^*+/-*^ mice with or without recent seizures (*Ldha, Ptpn5, Sergef*) (Fig. 2A-C), and two DEGs when comparing the strains within genotypes (*Ano5, Nav2*) (Fig. 2D-E). In terms of consequential coding sequence differences, we identified six genes in the refined *Dsm2b* interval with predicted missense or in-frame indel variants (*Dbx1, Myod1, Nav2, Prmt3, Sergef, Slc6a5*) (Table 2). Most missense variants were predicted to be tolerated or benign; however, variants in *Myod1* (rs13472315) and *Nav2* (rs32337268) were predicted to be probably damaging and possibly damaging/deleterious, respectively (Supplementary Table S1). Additionally, *Dbx1* has an in-frame deletion of 2 amino acids in 129 versus B6, and *Nav2* has an in-frame insertion that results in modest expansion of a polyglutamine repeat in 129. Among the nine genes with coding sequence or expression differences, six had a previous association with epilepsy and/or seizures based on literature and database searches (*Dbx1, Ldha, Nav2, Prmt3, Ptpn5, Slc6a5*) (Fig. 2F) (Table 2).

**Table 2.** *Dsm2b* Brain Expressed Candidate Genes.

| Gene | NCBI Gene ID | Location (GRCm39) | Expression Difference | Coding Difference (most severe consequence)* | Prior association with seizures and/or epilepsy |
| --- | --- | --- | --- | --- | --- |
| *Myod1* | 17927 | 7:46025898-46028523 |  | rs13472312 (m)<br>rs13472315 (m)<br>rs32790785 (m) |  |
| *Kcnc1* | 16502 | 7:46045921-46088128 |  |  | Yes |
| *Sergef* | 27414 | 7:46092578-46289231 | Yes | rs31674298 (m) |  |
| *Tph1* | 21990 | 7:46294065-46321961 |  |  |  |
| *Saal1* | 78935 | 7:46335532-46360104 |  |  |  |
| *Saa3* | 20210 | 7:46361422-46365124 |  |  | Yes |
| *Hps5* | 246694 | 7:46409890-46445488 |  |  |  |
| *Gtf2h1* | 14884 | 7:46445527-46473224 |  |  |  |
| *Ldha* | 16828 | 7:46490899-46505051 | Yes |  | Yes |
| *Ldhc* | 16833 | 7:46510627-46527566 |  |  |  |
| *Tsg101* | 22088 | 7:46538697-46569717 |  |  |  |
| *Uevld* | 54122 | 7:46572964-46608275 |  |  |  |
| *Misfa* | 629141 | 7:46633328-46637551 |  |  |  |
| *Spty2d1* | 101685 | 7:46640144-46658159 |  |  |  |
| *Tmem86a* | 67893 | 7:46700349-46704525 |  |  |  |
| *Ptpn5* | 19259 | 7:46727543-46783432 | Yes |  | Yes |
| *Zdhhc13* | 243983 | 7:48438751-48477188 |  |  |  |
| *Csrp3* | 13009 | 7:48480146-48497781 |  |  |  |
| *E2f8* | 108961 | 7:48516177-48531344 |  |  |  |
| *Nav2* | 78286 | 7:48558464-49259838 | Yes | rs250301556 (ins)<br>rs31226051 (m)<br>rs248206089 (m)<br>rs32337268 (m) | Yes |
| *Dbx1* | 13172 | 7:49281247-49286597 |  | rs246161289 (del) | Yes |
| *Htatip2* | 53415 | 7:49408863-49423723 |  |  |  |
| *Prmt3* | 71974 | 7:49428094-49508013 |  | rs32892158 (m) | Yes |
| *Slc6a5* | 104245 | 7:49559894-49613604 |  | rs31048165 (m) | Yes |
| *Nell1* | 338352 | 7:49624612-50516356 |  |  | Yes |
| *4933405O20Rik* | 243996 | 7:50248938-50250278 |  |  |  |
| *Ano5* | 233246 | 7:51160777-51248457 | Yes |  |  |
| *Slc17a6* | 140919 | 7:51271754-51320867 |  |  | Yes |
| *Fancf* | 100040608 | 7:51510325-51512015 |  |  |  |
| *Gas2* | 14453 | 7:51511763-51644723 |  |  |  |
| *Svip* | 75744 | 7:51646919-51655766 |  |  |  |
| *Ccdc179* | 100503036 | 7:51661431-51665476 |  |  |  |

\*Abbreviations: m, missense; ins, in-frame insertion; del, in-frame deletion

**Table 3.**
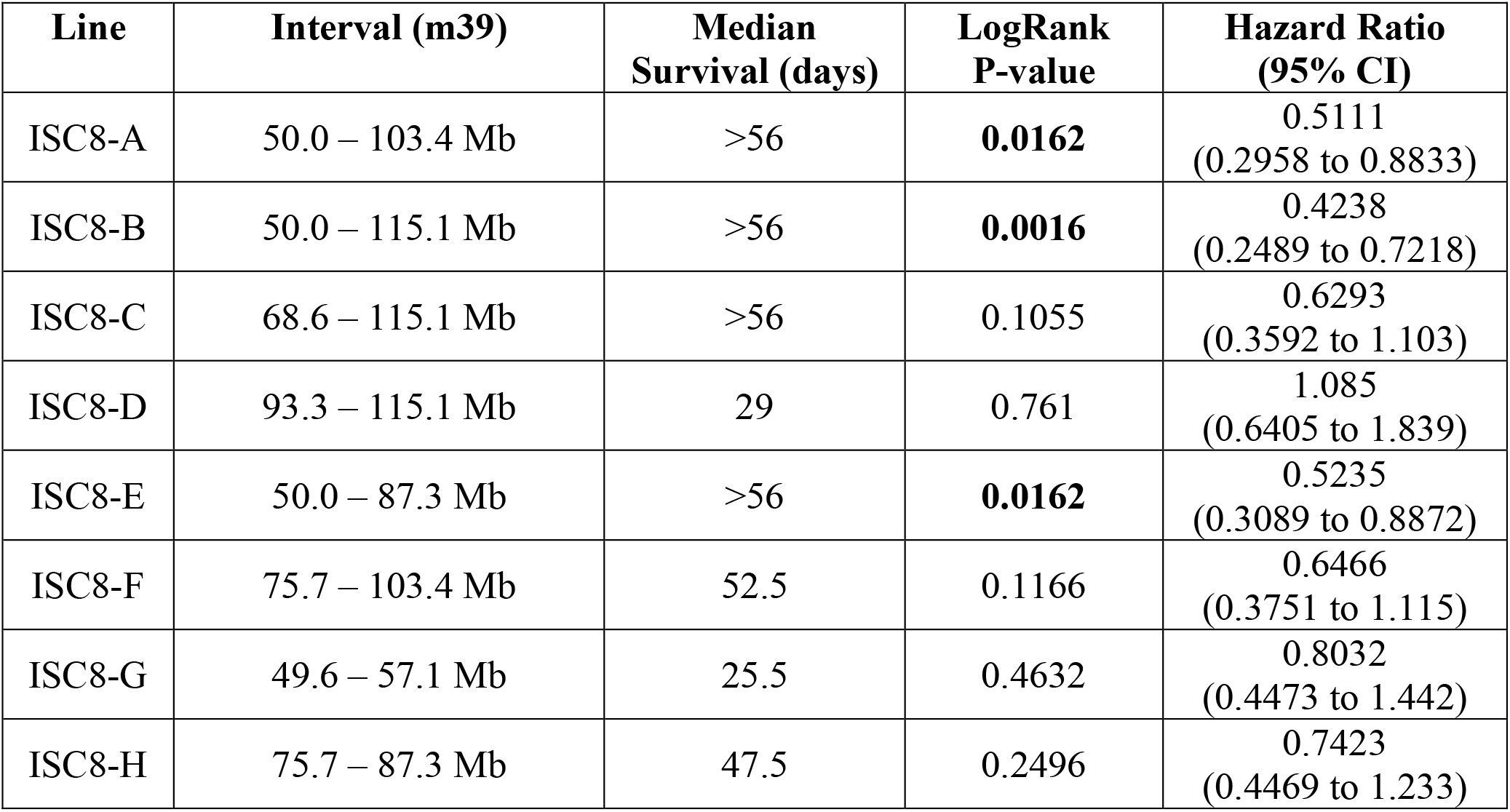
Interval specific congenic mapping of *Dsm3* on chromosome 8.

| Line | Interval (m39) | Median Survival (days) | LogRank P-value | Hazard Ratio (95% CI) |
| --- | --- | --- | --- | --- |
| ISC8-A | 50.0 – 103.4 Mb | >56 | **0.0162** | 0.5111<br>(0.2958 to 0.8833) |
| ISC8-B | 50.0 – 115.1 Mb | >56 | **0.0016** | 0.4238<br>(0.2489 to 0.7218) |
| ISC8-C | 68.6 – 115.1 Mb | >56 | 0.1055 | 0.6293<br>(0.3592 to 1.103) |
| ISC8-D | 93.3 – 115.1 Mb | 29 | 0.761 | 1.085<br>(0.6405 to 1.839) |
| ISC8-E | 50.0 – 87.3 Mb | >56 | **0.0162** | 0.5235<br>(0.3089 to 0.8872) |
| ISC8-F | 75.7 – 103.4 Mb | 52.5 | 0.1166 | 0.6466<br>(0.3751 to 1.115) |
| ISC8-G | 49.6 – 57.1 Mb | 25.5 | 0.4632 | 0.8032<br>(0.4473 to 1.442) |
| ISC8-H | 75.7 – 87.3 Mb | 47.5 | 0.2496 | 0.7423<br>(0.4469 to 1.233) |

**Table 4.**
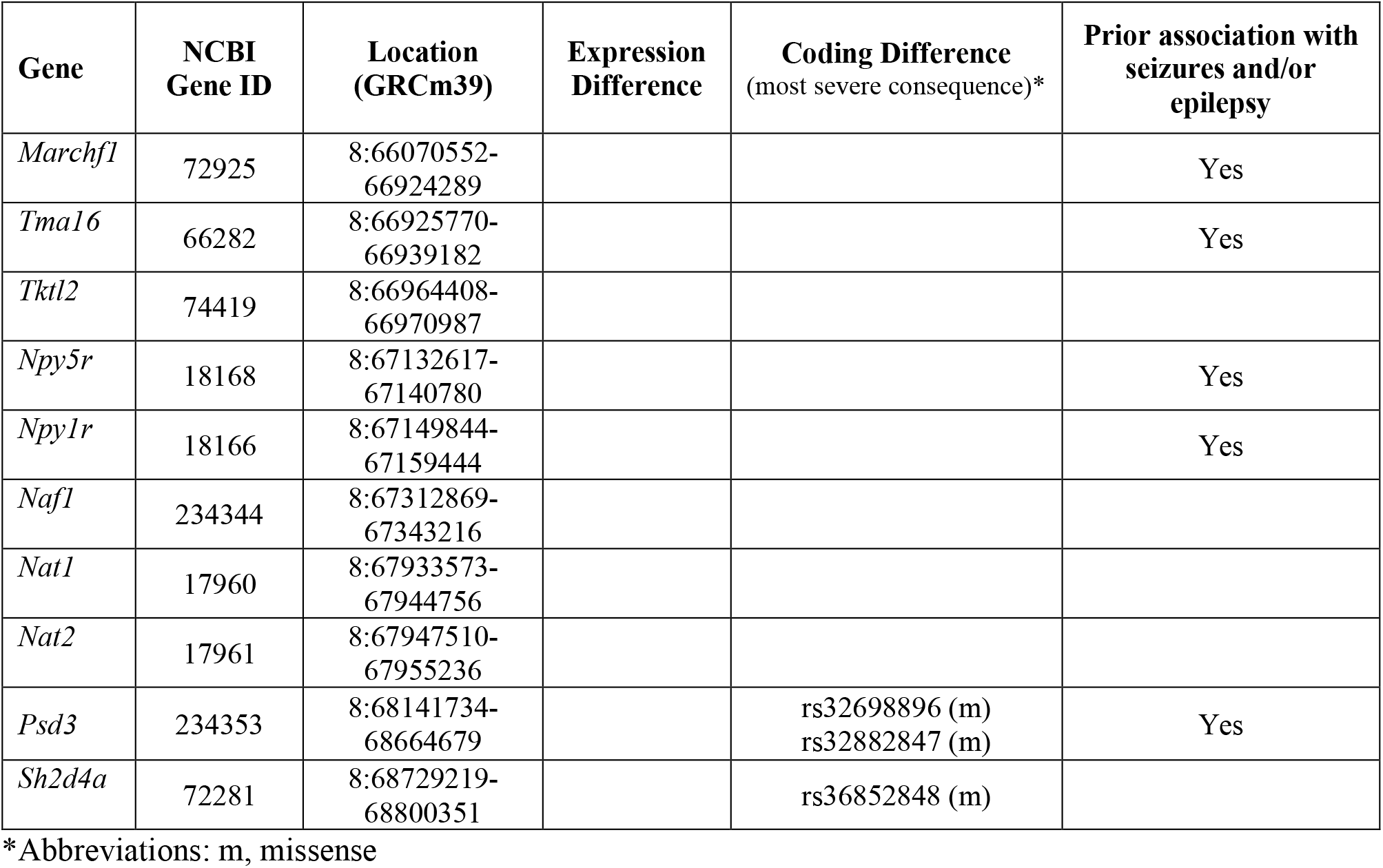
*Dsm3* Brain Expressed Candidate Genes.

**Figure 2.**
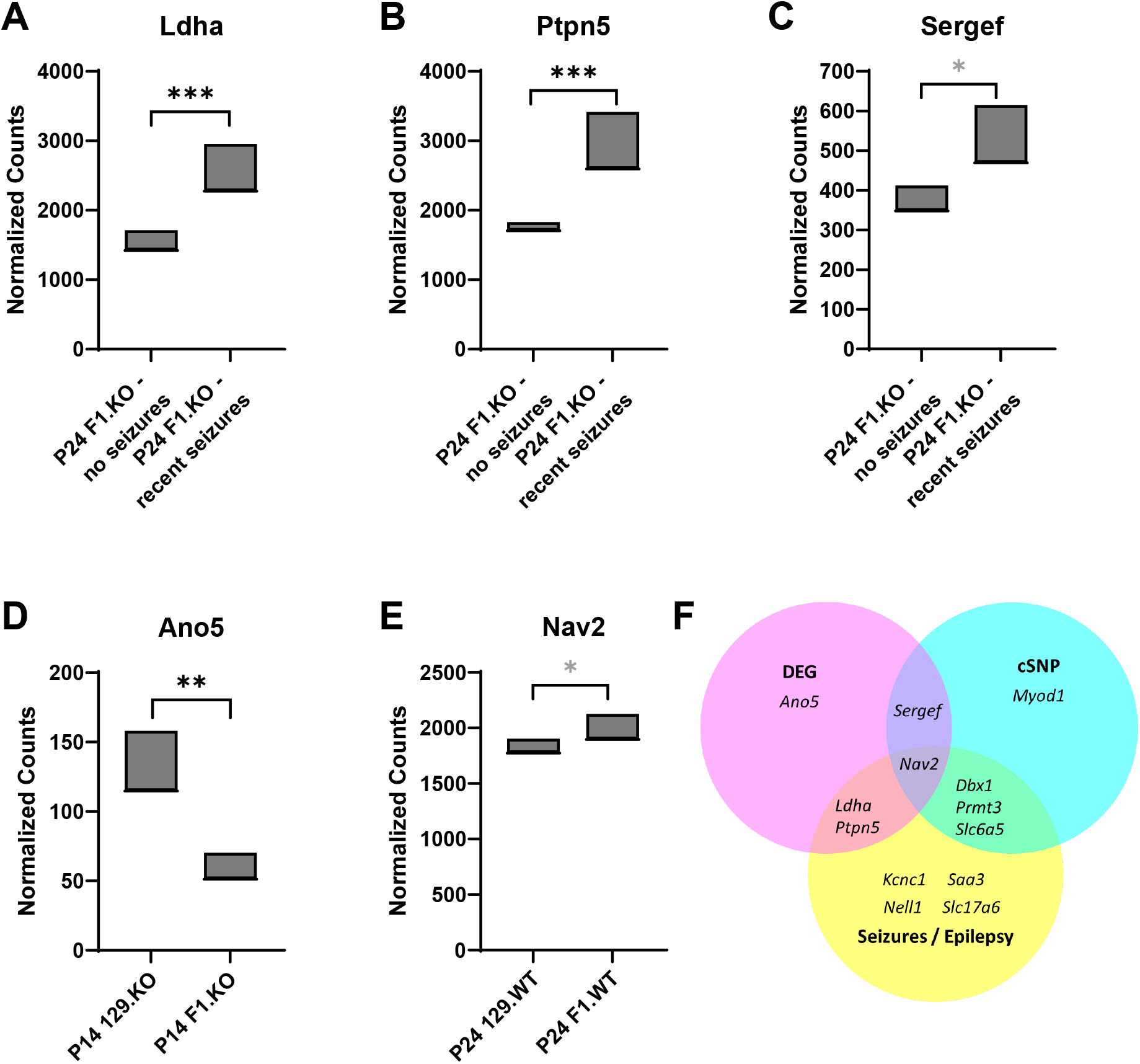
Analysis of *Dsm2* positional candidate genes. **A-E)** Differential expression of candidate modifier genes based on analysis of our previously published RNA-seq datasets (Hawkins et al. 2019; Hawkins et al. 2016). **A-C)** Comparing F1.*Scn1a*^*+/-*^ mice with or without seizures in the 24 hours preceding tissue collection, three genes were differentially expressed in hippocampus, *Ldha* (A), *Ptpn5* (B), and *Sergef* (C). **D)** *Ano5* had lower hippocampal expression in F1.*Scn1a*^*+/-*^ compared to 129.*Scn1a*^*+/-*^ mice at postnatal day 14 (P14). **E)** *Nav2* had lower hippocampal expression in wild-type (WT) 129 versus F1 mice at postnatal day 24 (P24). **F)** Summary of strain-dependent differences in candidate genes and their association with seizures and/or epilepsy. Six genes had strain-dependent differences and an existing biological link with epilepsy. ***FDR-adjusted *p*<0.0001, **FDR-adjusted *p*<0.003, *FDR-adjusted *p*<0.07 (from DeSeq2 reported in (Hawkins et al. 2019)).

### Dsm3 Fine mapping and Candidate Gene Analysis

For fine mapping of *Dsm3*, we used eight ISC lines with varying 129-derived intervals in the *Dsm3* region on chromosome 8 (Fig. 3A). Lines ISC8-B, ISC8-A and ISC8-E conferred improved survival, while other lines did not (Fig. 3B,C) (Table 2). Based on this, we localized the map interval to 66.2-68.8 Mb on chromosome 8.

**Figure 3.**
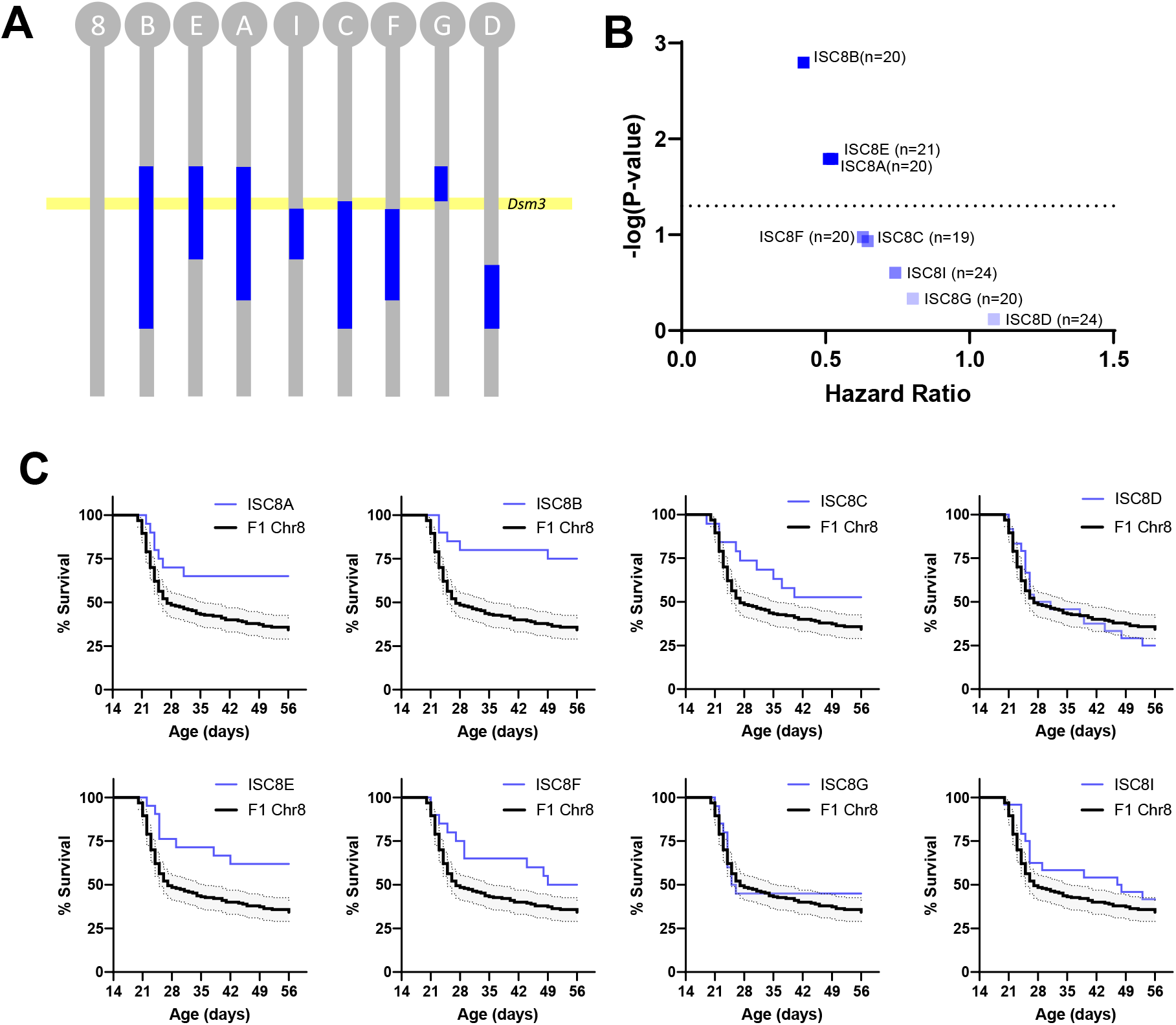
Fine mapping of *Dsm3* with ISC strains. **A)** *Dsm3* lines have varying 129-derived chromosome 8 intervals (blue) on a congenic B6 background (grey). B6.129-*Dsm3* ISC8 lines were crossed with 129.*Scn1a*^*+/*−^ mice and survival of *Scn1a*^*+/*−^ offspring was monitored to 8 weeks of age. **B)** Hazard ratios for all *Dsm3* ISC lines compared to F1 controls plotted against - log_10_(*P*-values) determined by Mantel-Cox LogRank test (n=19-24 per line). **C)** Kaplan Meier survival plots are shown for each ISC8 line (blue) compared to F1.*Scn1a*^*+/-*^ controls (black). Shaded area represents 95% confidence interval for F1.*Scn1a*^*+/-*^ controls.

The refined *Dsm3* interval contains 11 known protein coding genes. Of those, seven have confirmed expression in the brain. None of the brain expressed candidate genes have evidence of strain-dependent differences in gene expression, while two have missense coding sequence differences (*Psd3, Sh2d4a*) (Table 2). *Psd3*, encoding pleckstrin and Sec7 domain containing 3, has two missense variants between the strains that are each individually predicted to be tolerated/benign (Supplementary Table S1); however, they are in relatively close proximity and the combined effect of both substitutions is unknown. *Sh2d4a*, encoding SH2 domain containing 4A, has a missense variant that is predicted to be deleterious (Supplementary Table S1). Based on the literature and database searches, there is association with seizures or epilepsy for five of the brain expressed genes (*Marchf1, Tma16, Npy1r, Npy5r, Psd3*). Presently, *Psd3* is the only gene in the interval with evidence of strain-dependent differences and a biological association with seizures or epilepsy.

## DISCUSSION

In the current study, we used ISC mapping to refine two Dravet syndrome modifier loci, *Dsm2* on mouse chromosome 7 and *Dsm3* on chromosome 8. Chromosome 7 was complex, with additive contributions that suggested the contribution of at least two loci designated as *Dsm2a* and *Dsm2b*. Within the refined *Dsm2b* and *Dsm3* intervals we identified a number of modifier gene candidates that were prioritized based on (1) evidence of brain expression, (2) evidence of strain-dependent differences in expression or coding sequence, and (3) a plausible biological association with seizures/epilepsy. High priority candidates will be evaluated empirically in future studies to validate their modifier potential.

Among the *Dsm2b* genes, six were deemed strong candidates as they met our three levels of evidence. We further prioritized the genes based on strength of the biological association with seizures or epilepsy. *Nav2, Ptpn5, Ldha*, and *Dbx1* are deemed higher priority as they have a direct biological association with seizures or epilepsy, while *Prmt3 and Slc6a5* have an indirect association. For *Dsm3*, there was a single gene, *Psd3*, that met our three levels of evidence for a strong candidate. All strong candidates are discussed briefly below.

*Nav2* encodes Neuron navigator 2, a cytoskeletal associated protein important for cortical neuron migration and axon growth (Powers et al. 2022). Bi-allelic loss-of-function of *NAV2* was reported to cause a neurodevelopmental disorder with a complex brain malformation (Accogli et al. 2023). Disruption of the *Drosophila* ortholog (*Sickie*) results in hyperthermia-induced seizures and lethality (McNeill et al. 2011).

*Ptpn5* encodes Protein tyrosine phosphatase, non-receptor type 5, also known as Striatal-Enriched protein tyrosine Phosphatase (STEP). Deletion of *Ptpn5* in mice results in resistance to seizures induced by pilocarpine and attenuates audiogenic seizures in fragile X syndrome mice (Briggs et al. 2011; Chatterjee et al. 2018). In addition, pharmacological inhibition of STEP protects mice from seizures induced by kainic acid (Walters et al. 2022).

*Ldha* encodes lactate dehydrogenase A, a key enzyme in the astrocyte-neuron lactate shuttle. The lactate shuttle and lactate dehydrogenase (LDH) are inhibited by the ketogenic diet and stiripentol (Sada et al. 2015). Both are recommended therapies according to the Dravet syndrome International Consensus Panel, with stiripentol recommended as a second line treatment and ketogenic diet as a fourth line treatment (Wirrell et al. 2022). Furthermore, intrahippocampal antisense oligodeoxynucleotide knockdown of *Ldha* in mice suppressed high voltage spikes following kainic acid administration (Sada et al. 2015).

*Dbx1* encodes Developing brain homeobox 1, a transcription factor critical for interneuron differentiation, including Cajal Retzius cells in the cortex and brainstem pre-Bötzinger complex interneurons (Akins et al. 2017; Powers et al. 2022; Vann et al. 2018). Interestingly, the pre-Bötzinger complex is a critical nucleus for generating respiratory rhythms and has been implicated in SUDEP (Akins et al. 2017; George et al. 2023; Xia et al. 2016).

*Prmt3* encodes Protein arginine methyltransferase 3 that has been shown to methylate the cardiac NaV1.5 sodium channel and modulate its function (Beltran-Alvarez et al. 2013). Similarly, NaV1.2 function was modulated by *Prmt8*-mediated arginine methylation *in vitro*, while kainic acid-induced seizures *in vivo* resulted in elevated arginine methylation of NaV1.2 at three sites (Baek et al. 2014). Together, these observations raise the possibility of arginine methylation of NaV1.1 as a dynamic post-translation modification that could be mediated by *Prmt3*, which is expressed in brain.

*Slc6a5* encodes a presynaptic glycine transporter. Pathogenic variants in human *SLC6A5* result in hyperekplexia characterized by exaggerated startle response and life-threatening apneas that can result in sudden death in infancy (Rees et al. 2006). Although seizures are not a prominent feature of hyperekplexia, the strong association of *SLC6A5* with paroxysmal apneas suggests the possibility of enhanced SUDEP risk in the context of pre-existing epilepsy.

*Psd3* encodes pleckstrin and Sec7 domain containing 3, a guanine nucleotide exchange factor for ARF6 whose activity is critical for GABAergic synapse development (Kim et al. 2020; Sakagami et al. 2006). Knockdown of *Arf6* in mice results in enhanced seizure susceptibility, while dysregulation of ARF6 via *IQSEC2* pathogenic variants are associated with epilepsy, intellectual disability, and autism (Brant et al. 2021; Kim et al. 2020). Although we did not observe differential expression of *Psd3* in our transcriptomic datasets, a proteomics study of another Dravet mouse model (*Scn1a*^*A1873V*^) reported lower hippocampal expression of *Psd3* in Dravet versus WT mice at 4 weeks of age (Miljanovic et al. 2021).

The *Dsm2a* region is still large and requires additional refinement for efficient candidate gene analysis; however, it has not escaped our notice *Scn1b* is in this interval. *Scn1b* encodes the voltage-gated sodium channel beta subunit 1, which regulates NaV1.1 expression, localization, and channel gating (O’Malley and Isom 2015). It has already been reported that *Scn1a*^*+/-*^ mice exhibited a deficit in *Scn1b* expression, and that AAV-*Scn1b* administration targeting GABAergic neurons improved survival in *Scn1a*^*+/-*^ mice (Niibori et al. 2020). This provides support for *Scn1b* as a modifier of the *Scn1a*^*+/-*^ survival phenotype. However, because the mapped interval on proximal chromosome 7 is still large, the possibility of other contributing genes remains open.

Potential limitations of our study include the following. First, the *Dsm2* fine mapping data support the possibility of additive modifier genes on chromosome 7. This possibility could be empirically assessed with ISC lines that separate the proximal and distal intervals if the isolated effects sizes are large enough. However, in almost four years of breeding, recombination within this critical interval has not yet occurred in our colony that relies on natural recombination. It may be possible to engineer the desired recombination in the future. Recently, it was demonstrated that CRISPR/Cas9 editing can be utilized to circumvent recombination suppression in nematodes (Xie et al. 2023). Secondly, although we have evidence that some candidate genes are differentially expressed, absence of evidence should not be over-interpreted. Surveying expression of biologically plausible candidates across different brain regions and/or time points may reveal differences that were not captured in our existing datasets and could potentially elevate candidates that currently lack evidence of a strain-dependent difference. Similarly, absence of an existing association with epilepsy or seizures does not dismiss potential candidates; however, additional evidence is required to support a modifier role. Future studies may be necessary to gather additional evidence if high priority candidates fail to validate.

## Conclusion

Variable expressivity is common among individuals with Dravet syndrome due to *SCN1A* haploinsufficiency, supporting that genetic modifiers may contribute to clinical severity. It is challenging to discover modifier genes in humans in the context of rare disorders; therefore, we used the *Scn1a*^*+/-*^ Dravet mouse model as a tractable system for identification of modifier loci and candidate genes. Identifying modifier genes that influence clinical severity will advance our understanding of disease mechanisms and suggest potential targets for therapeutic intervention.

## DECLARATIONS

### Ethical Approval

Animal care and experimental procedures were approved by the Northwestern University Animal Care and Use Committee in accordance with the National Institutes of Health Guide for the Care and Use of Laboratory Animals.

## ACKNOWLEDGEMENTS

We thank Levi Barse, Tyler Thenstedt, Rylie Pancoast, Kelsey Davis, and Olivia Valente for technical assistance.

## DATA AVAILABILITY

Genomic and transcriptomic datasets analyzed during the current study are available in the NCBI GEO repository [GSE112627] and SRA repository [PRJNA817075]. Other datasets are available from [https://doi.org/10.18131/6qzh7-s1n82]

## CODE AVAILABILITY

Not applicable.

## AUTHOR CONTRIBUTIONS

Conceptualization: NAH, JAK

Formal analysis: JAK Investigation: NAH, NS

Methodology: NAH, JAK

Project administration: NAH, JAK

Supervision: NAH, JAK

Visualization: JAK

Writing – original draft: JAK

Writing – review & editing: NAH, JAK

Funding acquisition: JAK

