## Supplementary Table S1 for "Fine Mapping and Candidate Gene Analysis of Dravet Syndrome Modifier Loci on Mouse Chromosomes 7 and 8"

**Supplementary Table S1.** Effect of candidate gene coding variants determined by Ensembl Variant Effect Predictor (VEP).

| SYMBOL | Variant ID | Location | Consequence | IMPACT | Transcript (canonical) | cDNA position | CDS position | Protein position | Amino acids | Codons | Ref Allele | Alt Allele | SIFT Prediction (Score) | PolyPhen-2 Prediction (Score) |
|---|---|---|---|---|---|---|---|---|---|---|---|---|---|---|
| *Myod1* | rs13472312 | 7:46026253-46026253 | Missense | MODERATE | ENSMUST00000072514.3 | 356 | 157 | 53 | M/V | Atg/Gtg | A | G | Tolerated (1) | Benign (0.000) |
| | rs13472315 | 7:46026292-46026292 | missense | MODERATE | ENSMUST00000072514.3 | 395 | 196 | 66 | P/S | Cct/Tct | C | T | Tolerated (0.21) | Probably Damaging (0.997) |
| | rs32790785 | 7:46027237-46027237 | missense | MODERATE | ENSMUST00000072514.3 | 900 | 701 | 234 | A/V | gCg/gTg | C | T | Tolerated (0.06) | Benign (0.001) |
| *Sergef* | rs31674298 | 7:46092740-46092740 | missense | MODERATE | ENSMUST00000033127.12 | 1303 | 1268 | 423 | D/A | gAc/gCc | T | G | Tolerated_low_confidence (1) | Benign (0.000) |
| *Nav2* | rs250301556 | 7:49058454-49058454 | Inframe insertion | MODERATE | ENSMUST00000184945.8 | 673-674 | 576-577 | 192-193 | -/QQQQ | -/CAGCAACAGCAA | - | CAGCAA CAGCAA | - | - |
| | rs31226051 | 7:49114575-49114575 | missense | MODERATE | ENSMUST00000184945.8 | 2655 | 2558 | 853 | D/G | gAc/gGc | A | G | Tolerated_low_confidence (1) | Benign (0.000) |
| | rs32337268 | 7:49197728-49197728 | missense | MODERATE | ENSMUST00000184945.8 | 3453 | 3356 | 1119 | T/M | aCg/aTg | C | T | Deleterious_low_confidence (0) | Possibly damaging (0.578) |
| | rs248206089 | 7:49197731-49197731 | missense | MODERATE | ENSMUST00000184945.8 | 3456 | 3359 | 1120 | V/A | gTc/gCc | T | C | Tolerated_low_confidence (1) | Benign (0.000) |
| *Dbx1* | rs246161289 | 7:49282284-49282290 | Inframe deletion | MODERATE | ENSMUST00000032717.7 | 1048-1053 | 914-919 | 305-307 | PAH/H | cCGGCGCac/cac | GCGCCG | - | - | - |
| *Prmt3* | rs32892158 | 7:49448109-49448109 | missense | MODERATE | ENSMUST00000032715.13 | 949 | 818 | 273 | V/A | gTt/gCt | T | C | Tolerated (1) | Benign (0.000) |
| *Slc6a5* | rs31048165 | 7:49561578-49561578 | missense | MODERATE | ENSMUST00000056442.12 | 366 | 109 | 37 | T/A | Acg/Gcg | A | G | Tolerated_low_confidence (1) | Benign (0.000) |
| *Psd3* | rs32698896 | 8:68573659-68573659 | missense | MODERATE | ENSMUST00000212960.2 | 1131 | 521 | 174 | G/E | gGg/gAg | C | T | Tolerated_low_confidence (0.17) | Benign (0.000) |
| | rs32882847 | 8:68573714-68573714 | missense | MODERATE | ENSMUST00000212960.2 | 1076 | 466 | 156 | R/G | Aga/Gga | T | C | Tolerated_low_confidence (1) | Benign (0.000) |
| *Sh2d4a* | rs36852848 | 8:68787743-68787743 | missense | MODERATE | ENSMUST00000066594.4 | 1352 | 848 | 283 | I/T | aTc/aCc | T | C | Deleterious (0.01) | Benign (0.000) |
